# Rapid thermal adaptation in coral photosymbionts draws on standing variation and recombination

**DOI:** 10.64898/2026.09.08.749829

**Authors:** Patrick Buerger, Heng L. Yeap, Owain R. Edwards, Madeleine J.H. van Oppen, John G. Oakeshott

## Abstract

Bleaching tolerance in corals depends in part on the thermal tolerance of their microalgal symbionts. Laboratory evolution has increased the thermal tolerance of the symbiont *Cladocopium proliferum* in ∼120 generations, but the genetic basis of that response was unknown. We compared single nucleotide polymorphisms in transcriptomes of three heat-evolved *C. proliferum* strains and one wild-type (unselected) strain from the same progenitor. We found 15,640 polymorphic loci, but no variant was both private to a strain and consistent across its replicates, which indicates that new mutations contributed little to the response in expressed sequences. Instead, allele frequencies at 350 loci differed significantly between strains, and linkage patterns indicated recombination had occurred within scaffolds both before and after the strains were separated. Selection and recombination of variation already present in the progenitor therefore underpinned the rapid thermal adaptation. Experimental evolution for reef restoration should start from genetically diverse cultures rather than single cell isolates.

## INTRODUCTION

Many coral reefs worldwide have declined due to mass mortalities over the last few decades caused by the increasing frequency and severity of marine heatwaves (1, 2). A reason for the coral’s temperature sensitivity is their symbiotic relationship with microalgal symbionts (Symbiodiniaceae), which can break down when seawater temperatures exceed long-term average summer maxima, resulting in the loss of the algal cells from the coral tissues (i.e., coral bleaching) and potential mortality (3). Both partners contribute to the holobiont’s overall bleaching tolerance (4–6) but unraveling the underlying genetic and molecular mechanisms in each has proven difficult. The contribution of the coral host to tolerance is a polygenic trait determined by the combined effects of numerous genes and their interactions (7, 8). For Symbiodiniaceae, transcriptomic comparisons across strains and species show that the same thermal stress can produce different gene-expression responses (9, 10), which indicates genotype-dependent variation in response but also suggests that interactions among genes can contribute to differences in thermal tolerance. In addition, it has proven difficult to develop high-quality reference genome annotations for Symbiodiniaceae, which have unusually large numbers of genes that lack homologues in other, better characterized taxa. Thus, high proportions of their genes are still functionally anonymous (11–14).

One promising approach to the study of thermal tolerance mechanisms in the microalgae involves experimental evolution (15, 16). This approach exploits the short generation times of Symbiodiniaceae and the abilities to culture them in the laboratory, subculture them into presumptively clonal cell lines and propagate those lines under different thermal regimes. Experimental evolution has produced strains of *Cladocopium proliferum* with enhanced *in vitro* thermal tolerance in just 80 generations over 2.5 years (17), and comparable responses in other Symbiodiniaceae species after ∼50 generations (16, 18). Moreover, assays of later generations showed a subset of the *C. proliferum* heat-evolved strains also conferred their thermal tolerance into the symbiosis with coral: *Acropora kenti* (formerly *A. tenuis*) larvae (19), *A. kenti* juveniles (20), *A. spathulata* recruits (21, 22) and adult *Galaxea fascicularis* (23). Heat-evolved strains had enhanced ROS scavenging capacities via elevated levels of reduced glutathione (24) and/or produced lower levels of ROS, as indicated by reduced amounts of extracellular ROS (17, 19) *in vitro*. At least one tolerance-conferring strain exhibited enhanced protection of its photosystems against oxidative stress through higher levels of carotenoid-like pigments (23), while transcriptomics suggested it also had relatively lower rates of photosynthesis and increased expression of carbon fixation genes *in hospite* compared to the other heat-evolved strains (19).

Building on these findings, we analyzed single nucleotide polymorphisms (SNPs) in three heat-evolved *C. proliferum* strains and a wild-type strain derived from the same progenitor stock, using published transcriptomes (19), and tested for associations between these variants and their thermal tolerance phenotypes. Thermal adaptation drew largely on standing variation in the progenitor rather than on new mutations, and recombination reorganized that variation into new haplotype combinations both before and during selection. This has a direct consequence for reef restoration, insomuch as experimental evolution should combine multiple genotypes and not start from single cell isolates, so that selection has existing variation to act on instead of relying on new mutation.

## MATERIALS AND METHODS

### Experimental Design

The progenitor *Cladocopium* strain (formerly *Cladocopium* C1^acro^ (62, 63) was isolated from an *A. kenti* coral collected from the inshore, central Great Barrier Reef (GBR) in 2010 (17). The extracted cells were first maintained as a heterogeneous culture, from which a clonal strain was later established by picking cells from a single colony forming unit on solid phase media, which was assumed to produce a clonal culture (17). Ten replicate subcultures were then subjected to a step-wise temperature increase from 27°C to 31°C and kept at 31°C (17). After 4 years (∼120 generations), all ten heat-evolved strains showed more thermal resistance than wild-type control subcultures from the same progenitor stock in culture, while only three conferred tolerance in symbiosis with *A. kenti* larvae under thermal stress (one-week exposure to 31°C) (19).

We analyzed transcriptomes generated previously from four of these strains in symbiosis with *A. kenti* larvae at ambient temperature (27°C) (19): The heat-evolved strain conferring thermal tolerance (SS8), two heat-evolved strains that did not (SS3 and SS5), and one wild-type control strain (WT10). Ten larvae were pooled into each experimental replicate, with six replicates for each of SS3, SS5 and WT10, and five for SS8. The raw data is available at NCBI Gene Expression Omnibus (GEO), repository GSE133082, BioProject PRJNA549921: SRA SRP201999. Full culture, selection, sequencing and coverage details are given in Supplementary Methods S1 and S2.

### SNP analyses

SNP calling was performed using the Genome Analysis Toolkit best practices workflow (GATK version 4.3.0.0, “RNAseq short variant discovery (SNPs + Indels)” at https://gatk.broadinstitute.org/hc/en-us/articles/360035531192, (64). As recommended, RNAseq reads were mapped with STAR aligner (version 2.5.3a, (65)) in 2-pass mode to the *C. proliferum* v1.0 reference genome (25, 66) in order to separate host from Symbiodiniaceae reads. Variants were filtered on read depth, strand bias, variant confidence and sequencing bias following previous population genomic studies of coral holobionts (67, 68), and singleton variants and loci without sufficient read coverage in all samples were removed. Software versions, filter thresholds and command line parameters are listed in Supplementary Methods S3. Triallelic loci were split into biallelic loci (bcftools, v1.10.2). SNP impacts on ORFs were checked with SNPeff (v5.0) using default settings and classified as up- or downstream if they were located within 1,000 nt distance of an ORF. Functional analyses of the SNPs were based on published gene predictions for the reference *C. proliferum* genome (25) and the annotations of a previous RNA-seq analysis (19).

SNP patterns among the four strains were first compared by principal component analysis (PCA, *mixomics* R package; (69)). One SS8 replicate (405S14) with relatively low sequence depth was identified as an outlier (Fig. S1), excluded, and the PCA rerun. Pairwise Fst distances among strains were calculated with *poolfstat* (70) and used to generate a phylogenetic dendrogram with *phylip* (version 3.698; (71)), visualized with *FigTree* (version 1.4.4; github.com/rambaut/figtree/).

Significant intergenic SNPs were screened for previously unannotated features by analyzing their genomic regions for novel ORFs (≥500 nt) using Artemis software (72). Newly detected ORFs were functionally annotated via BLASTp (non-redundant database, updated: 2025/05/28) (73) with a match threshold of ≥60% query coverage and an e-value of ≤ 1 x 10-20. Overlaps of significant intergenic SNPs with loci encoding long non-coding RNAs (lncRNAs) were also identified after mapping previously identified lncRNAs from Cladocopium (26) to its reference genome using BLASTn (threshold e-value: <1e-05; sequence identify: 100%). SNP-lncRNA associations were cross-referenced with modules of a previously defined thermal-tolerance weighted gene co-expression network analysis (WGCNA) (26).

*ClusterProfiler* (version 4.4; (74)) was used to check for enriched Molecular Function (MF) and Biological Process (BP) Gene Ontology (GO) categories among significant SNPs with functional annotations using a generic GO-slim list (goslim_generic.obo) and an FDR-corrected significance threshold of p-adjust < 0.01. The minimum gene set size (minGSSize) for the GO analyses was set to 1 to ensure generally rare categories were still considered.

*LDx* software (75) for pooled sequencing data was used to obtain maximum likelihood estimates (mle_est) of linkage disequilibrium (LD) between pairs of SNPs in each scaffold containing at least six SNPs, and to model the relationship of mle_est to the distance separating the respective SNPs in each of those scaffolds (76) (Supplementary Methods S4). Models could be obtained for 129 of the 711 scaffolds.

### Statistical Analysis

Allele frequency differences between strains were plotted in R using *ggplot2* (version 3.5.2; (77)), and tested for significance using linear modelling and analysis of variance, with a significance threshold for p-adjust of 0.01 after Bonferroni correction. Significant differences were presented using the R package *emmeans* (v1.10.5; (78)). Three contrasts were used to assess the relationships of the significant SNPs to the thermal properties of the respective strains: one between WT and the three SS strains to assess their roles in *in vitro* tolerance, and two between SS8 and SS3 or SS5 to assess their roles in holobiont tolerance. Chi-squared tests (χ^2) with Bonferroni corrections for post-hoc comparisons (significance threshold for p-adjust of 0.01) were applied using the R package *stats* (version 3.6.2; (79)) to compare the distribution of the significant SNPs across impact categories with that of the non-significant SNPs.

In order to understand the effects of linkage and recombination on patterns of allele frequency differences between strains the SNP data for each scaffold containing at least six SNPs (n = 711) was subjected to a two-way analysis of variance (ANOVA: Python package *statsmodels* 0.13.5 (80)) testing for main and interaction effects of strain and SNP. Significance thresholds after Bonferroni correction were again set at p-adjust < 0.01, but two additional filters of R-squared ≥ 0.20 and minimum allele frequency difference of ≥ 0.20 were also used to focus on the larger effects.

## RESULTS

### SNP profiles

SNPs were called from published transcriptomes of *A. kenti* larvae hosting four *C. proliferum* strains: three heat-evolved strains (SS3, SS5 and SS8, of which only SS8 conferred thermal tolerance on the host) and the unselected wild-type strain WT10. After filtering and quality control 15,640 bi-allelic SNPs were retained (15,564 bi-allelic and 38 tri-allelic SNPs split into 76 bi-allelic SNPs).

The SNPs were distributed across ∼11% (4,739 of 41,289) of the scaffolds in the reference assembly, which represented 36% (431 Mbp of 1.2 Gbp) of the assembly’s length and covered 14% (4,884 of ∼35,000) of its annotated protein-coding genes (25). Scaffolds with at least three SNPs accounted for 75% of all the SNPs and one scaffold, #4420 (66 kbp long, four genes), contained 89 SNPs (Table S1). Without a chromosome-level reference assembly, the linkage relationships among the scaffolds containing the SNPs could not be determined.

A principal component analysis of allele frequencies for all 15,640 biallelic SNPs found that PC1, explaining 19% of the variance, separated WT10 from the three heat-evolved strains (Fig. 1A). Fst distances between the four strains (Fig. 1B, 1C) recapitulated this difference. Although the four SS8 replicates (conferring tolerance) retained after exclusion of a low coverage outlier clustered together in both the PCA and the Fst dendrogram, they showed only minor differences from SS3 and SS5 (not conferring tolerance) in either the PCA or Fst analyses.

**Fig. 1.**
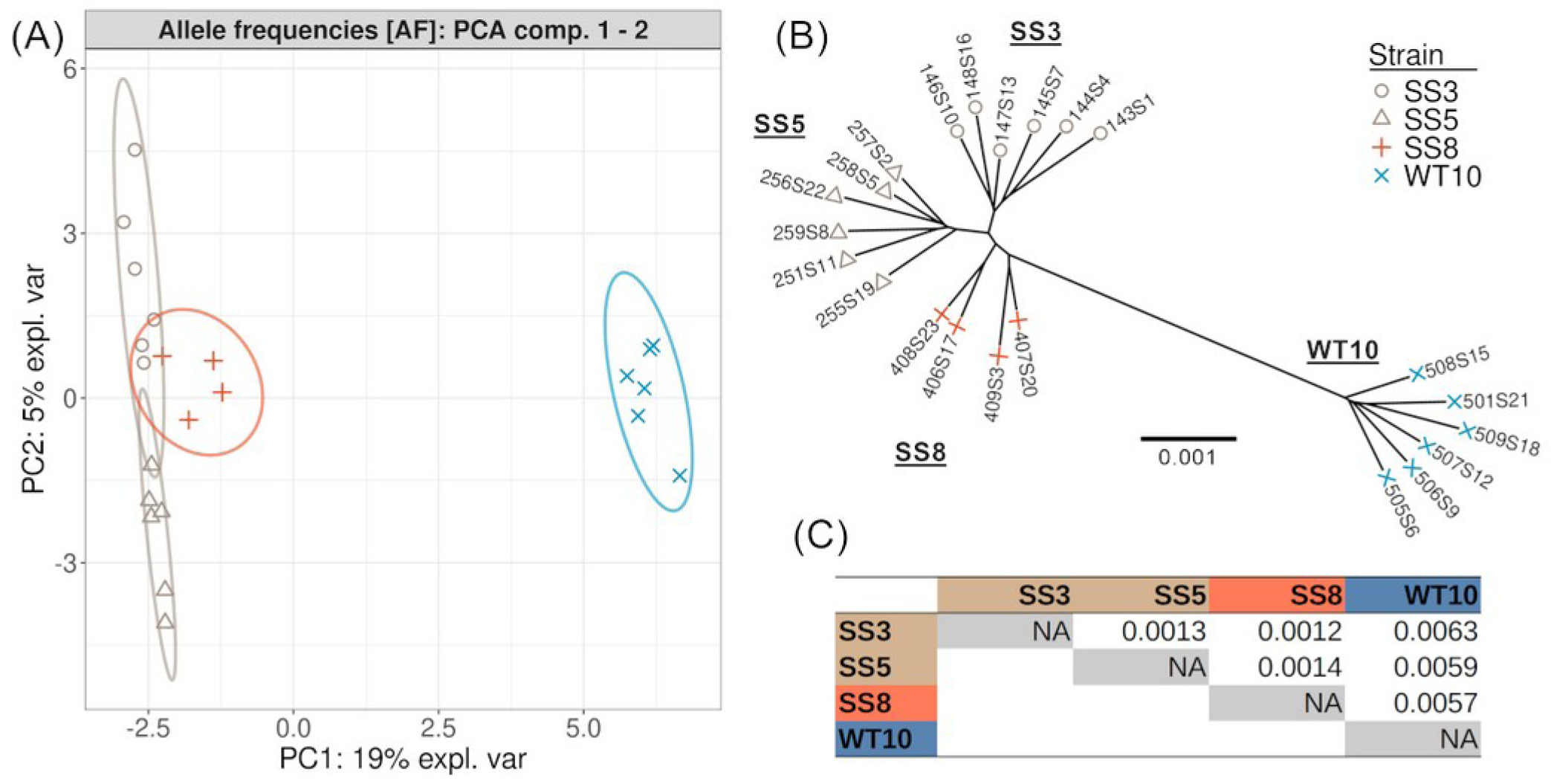
Genetic relationships among strains (15,640 SNPs). (A) PCA of the allele frequency data. (B) Dendrogram based on Fst distances between samples. (C) Pairwise Fst values between the strains (Fst-group).

The great majority of the SNPs were found to be polymorphic in all four strains and fewer than 1% of them were confined to all replicates of a single strain (Fig. 2A, S2A). However, linear modelling and analyses of variance found significant frequency differences among strains for 350 (∼2%) SNPs. These significant SNPs were distributed across 155 scaffolds that covered 16 Mbp (or 1.3%) of the genome (Table S2). Two hundred and thirteen were polymorphic across all replicates of all four strains, 46 were fixed in some replicates of a strain, and 91 reached fixation in all replicates of at least one strain (Fig. 2A, 2B), while none were private for a particular strain (Fig. S2B). Ninety of the 91 fixed in all replicates of at least one strain were fixed in SS3 and none were fixed in WT10. Allele frequencies for most of the 350 significant SNPs were at intermediate values in WT10, whereas many showed significant shifts towards more extreme values in SS3, SS5 and SS8 (Fig. 2B; χ^2 p-values < 0.0001, Table S3).

**Fig. 2.**
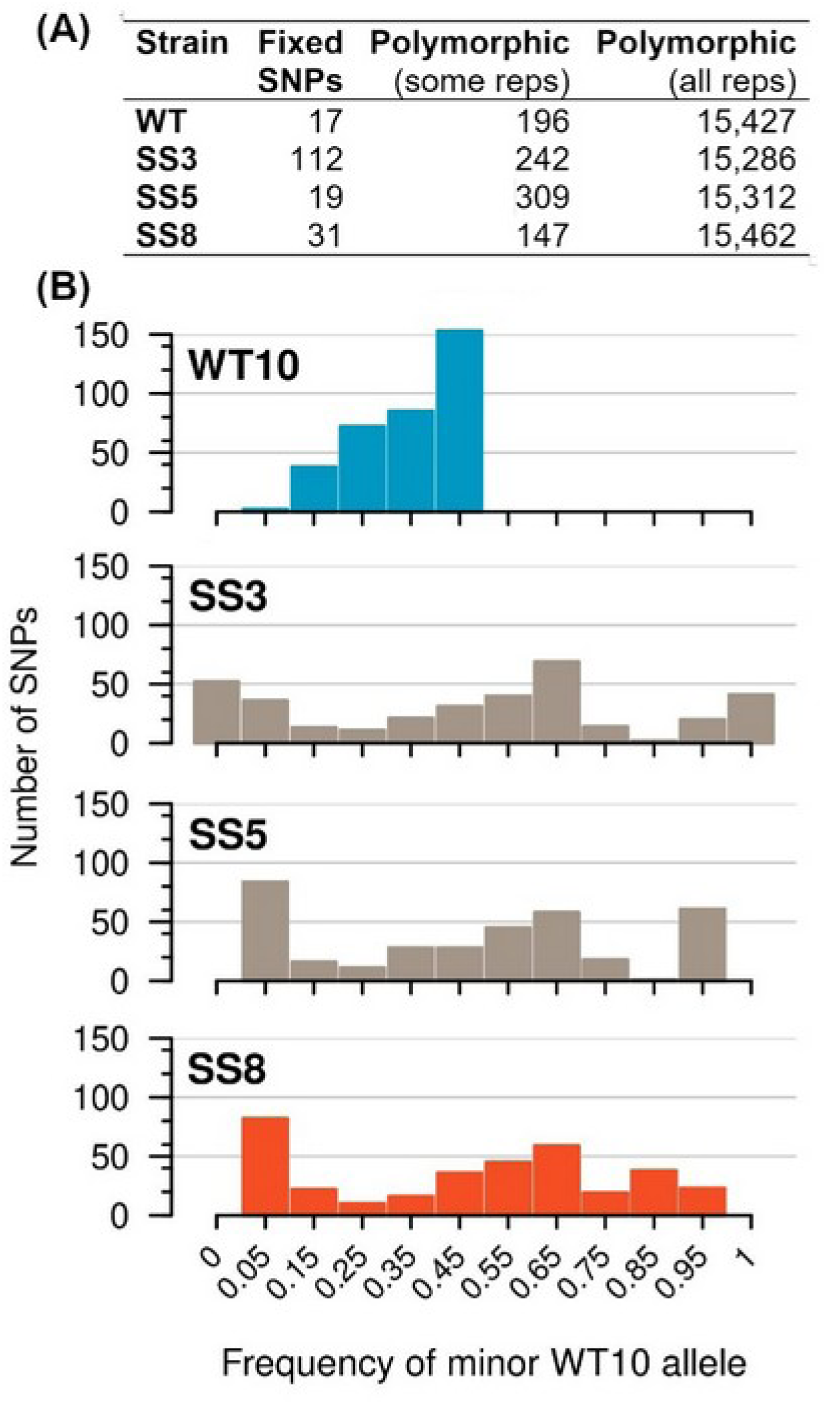
Frequency distribution of single nucleotide polymorphism within strains. (A) The total numbers of SNPs in each strain (15,640 loci), categorized as either fixed in all replicates or polymorphic in some or all replicates. (B) Frequency histograms of the 350 significant SNPs in each strain for the minor allele in WT10. While many of these SNPs had intermediate allele frequencies in WT10, they were closer to fixation in the SS strains, especially SS3 and SS5. All pairwise comparisons of the four distributions in (B) were significantly different, except the one between SS5 and SS8 (Table S3).

### WT10 vs SS strains

Consistent with the PCA results, the great majority, 339, of the 350 significant SNPs differentiated the three SS strains from WT10. The distribution of these 339 SNPs across functional impact categories was broadly similar to that of the non-significant SNPs (Table 1) and, given that the SNPs were sourced from transcriptome data, the proportions of intergenic variants in both groups were surprisingly high (∼31% and 33%, respectively). However, manual annotation revealed that 18 of the 115 significant intergenic SNPs lay in 7 different, previously unannotated ORFs, while 32 lay in 18 lncRNA genes (Supplementary Material S1) when mapped against a lncRNA database (26).

**Table 1.** Summary of predicted impacts of all SNPs on coding and non-coding regions. The numbers of SNPs in each impact category are shown in the ‘#’ columns, along with their percentage of the total SNPs analyzed in the % columns. Both significant and non-significant SNPs were distributed relatively evenly between coding and non-coding regions, with just slightly lower proportions in flanking variants in the latter (χ^2 = 14.4, p ∼0.03).

| SNP impact | region | Non-significant SNPs |  | Significant SNPs |  |  |
| --- | --- | --- | --- | --- | --- | --- |
|  |  | # | % | WT vs | SS8 vs | % |
|  |  |  |  | SS | SS3, SS5 |  |
| Non-synonymous | genic | 3,757 | 24.6 | 93 | 1 | 26.9 |
| Disruptive / Frameshifts | genic | 525 | 3.4 | 5 | 3 | 2.3 |
| Synonymous | genic | 3,460 | 22.6 | 62 | 2 | 18.3 |
| Intron | genic | 1,134 | 7.4 | 19 | 0 | 5.4 |
| Downstream | flanking | 1,024 | 6.7 | 29 | 0 | 8.3 |
| Upstream | flanking | 599 | 4.0 | 21 | 0 | 6.0 |
| Intergenic region | intergenic | 4,791 | 31.3 | 110 | 5 | 32.9 |
| **Sum (total of 15,640 SNPs)** |  | **15,290** | **100%** |  | **350** | **100%** |

Two hundred and thirty five of the significant SNPs were categorized as having some impact (Table 1) on a total of 130 unique genes (Supplementary Material S1), and their functional annotations suggest some of these genes could be involved in heat stress responses and/or enhanced thermal tolerance (Table S4). Notable examples included a putative ubiquitin protein ligase HERC2, a carbonic anhydrase 2, a 9-cis-epoxycarotenoid dioxygenase, a peroxisome membrane protein, peroxisome biogenesis factor, light-harvesting protein, and a glutathione S-transferase. The three GO categories for genes that were significantly enriched among the significant SNPs as compared to those in the total SNP dataset were regulator activity (MF, 5x; GO:0098772; p-adjust < 0.005), carbohydrate metabolic process (BP, 11x; GO:0005975; p-adjust < 0.01) and DNA replication (BP, 5x; GO:0006260; p-adjust < 0.0001).

The highest number of significant SNPs in, or flanking, an individual gene was in a gene with a non-ribosomal peptide synthetase terminal domain of unknown function (NRPS_term_dom, gene ID: SymbC1.scaffold12047.2.m1). Twenty three of the 28 NRPS_term_dom SNPs differed significantly between the SS strains and WT and, of these 23, 4 were missense, 1 synonymous, 17 downstream variants and 1 upstream. There were also relatively high numbers of significant SNPs, 18 in total, in four genes coding for ubiquitin protein ligases, namely HERC1, HERC2, HERC4 and ARI5 (scaffold513.gene1 manually annotated, scaffold1827.1.m1, scaffold2331.2.m1, and scaffold5652.2.m1). These 18 differences included 11 missense, 2 synonymous, 2 upstream and 2 downstream variants, as well as 1 disabling frameshift.

Nine of the 130 previously annotated genes that were impacted by significant SNPs had also previously been found to differ significantly in expression levels (>0.5 lfc, <0.05 FDR) between larvae hosting SS vs WT strains (19). For example, the NRPS_term_dom gene above and a gene encoding a light-harvesting protein (scaffold75.5.m1) were significantly downregulated in the SS as opposed to WT10 strains.

All lncRNA genes that contained significant SNPs in the WT10 vs SS comparisons were also found to be associated with regulatory modules in the Weighted Gene Co-expression Network Analyses (WGCNA) (26). In particular, one lncRNA gene (Cproliferum_lnc.39531, on scaffold1429) which contained 10 of the 33 significant SNPs located in lncRNAs was found in WGCNA Module 22, whose expression showed the strongest positive expression correlation with temperature (+0.957) in previous heat stress experiments (26).

### SS8 vs SS3 and SS5

Allele frequencies in just 11 SNPs across five scaffolds (1282, 3829, 6722, 8952 and 14903, with scaffold 3829 also present in the WT10 versus SS comparison) differed between SS8 and/or SS3 and SS5. None of these SNPs were also significant in the SS vs WT contrasts above and none showed large differences between SS8 and the other two SS strains. Three of the 11 SNPs were in confirmed intergenic regions, and another one in a gene for an unknown protein.

However, seven of the 11 SNPs lying in just two scaffolds, 3829 and 8952, were of particular interest. Scaffold 3829 contained three SNPs, namely a synonymous difference in a carbonic anhydrase 2 gene (scaffold3829.2.m1) and two intergenic variants between this and another carbonic anhydrase gene. Neither of the two carbonic anhydrase genes had been found to be differentially expressed between SS8 and the other two SS strains (19). Scaffold 8952 contained four of the 11 SNPs, three of them frameshifts and one synonymous in a 2132 nt ORF which blast searches revealed had similarities to an amino acid transporter (AVT1J, *Symbiodinium pilosum*, query coverage >56%, e-value <1*10-80).

### Recombination and linkage disequilibrium

Allele frequencies within each of the 711 scaffolds carrying at least six SNPs were tested for effects of strain, SNP identity and their interaction with two-way ANOVAs, and linkage disequilibrium (LD) between SNP pairs was estimated across the same set of scaffolds.

The ANOVAs for 268 of the 711 scaffolds found no large main or interaction effects for SNP or strain differences on allele frequencies (Table 2A, Fig. 3A, Supplementary Material S2), which suggests these were parts of pre-existing haplotypes unaltered by recombination and unaffected by the thermal selection.

**Fig. 3.**
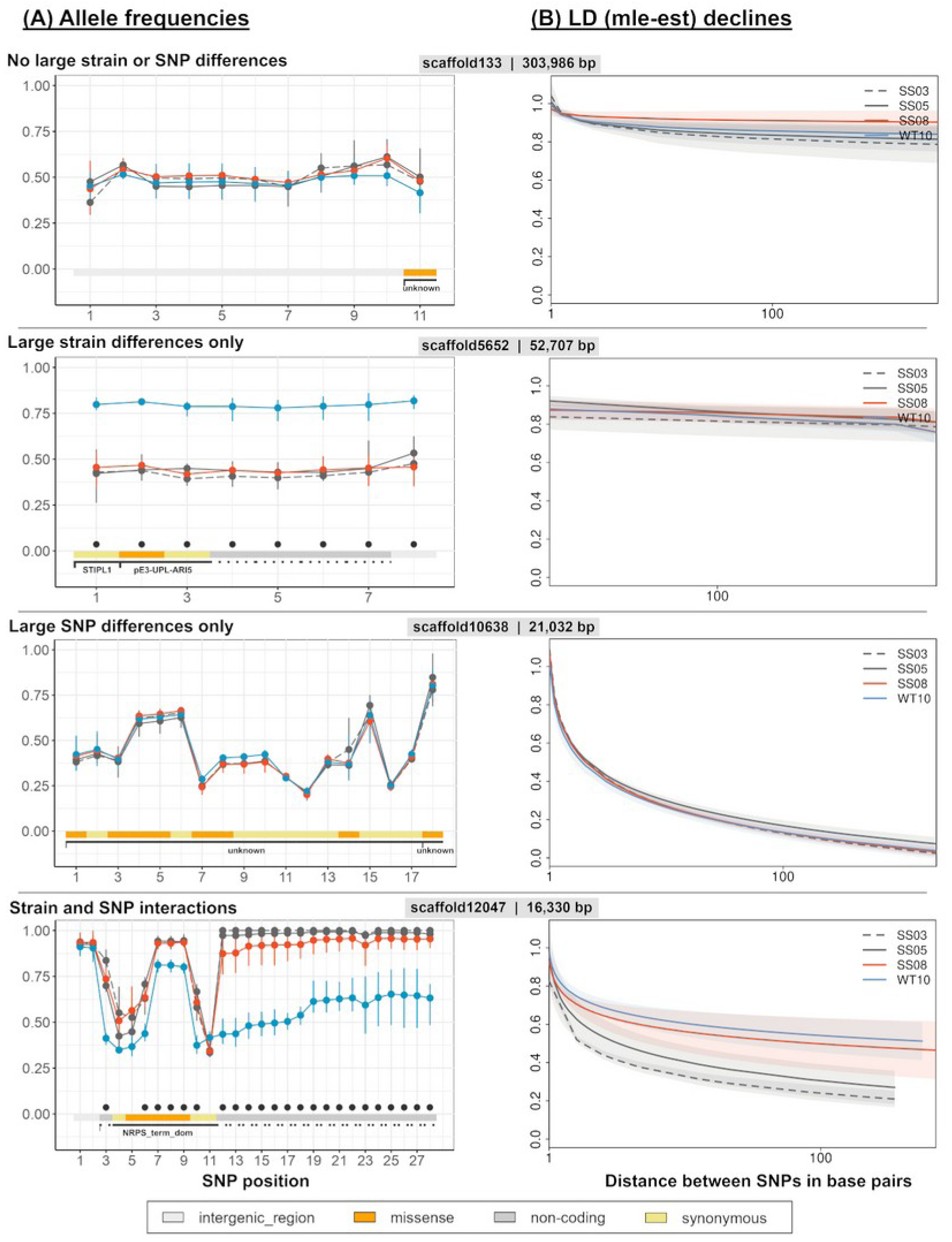
Examples of the four major patterns of large intra-scaffold (A) allele frequency differences and (B) linkage disequilibrium as maximum likelihood estimates (mle_est). (A) SNPs are numbered according to their order on the scaffolds. Black dots at the bottom of a plot indicate significant differences between strains, while the colored bars indicate SNPs with potential functional impacts on genes. Gene names and their SNP coverage are abbreviated below the colored bars, with the broken lines indicating up-/downstream regions of a gene, solid lines represent genes with their potential function (unknown: unknown protein; STIPL1: Septin and tuftelin-interacting protein 1-like 1; pE3-UPL-ARI5: putative E3 ubiquitin-protein ligase ARI5; NRPS_term_dom: non-ribosomal peptide synthetase terminal domain of unknown function). (B) Lines show the mean fitted linkage disequilibrium (maximum likelihood estimate) as a function of distance between SNP pairs, averaged across replicates within each strain, while shaded regions lie within a standard deviation of the mean mle_est.

**Table 2.** Summary of the ANOVAs on allele frequencies (A) and LD (B) results. Only the 711 scaffolds with at least six SNPs were considered for the ANOVAs on allele frequencies, and only 129 of these (generally those with the most SNPs) generated maximum likelihood estimates (mle_est) of linkage disequilibrium. The 711 scaffolds contained 165 of the significant SNPs.

| (A) | Category | Large effects |  |  | Number of scaffolds (n = 711) | Significant SNPs |  |
| --- | --- | --- | --- | --- | --- | --- | --- |
|  |  | SNP | Strain | SNP x strain |  | WT vs. SS | SS8 vs. SS3, SS5 |
|  | None | - | - | - | 268 (38%) | 8 | 0 |
|  | Strain only | - | Y | - | 8 (1%) | 54 | 0 |
|  | SNP only | Y | - | - | 360 (51%) | 10 | 0 |
|  | SNP x strain | - | - | Y | 13 (2%) | 4 | 0 |
|  |  | Y | - | Y | 58 (8%) | 29 | 11 |
|  |  | Y | Y | Y | 4 (1%) | 49 | 0 |
| (B) | Category | Number of scaffolds (n = 129) |  | mle-est [mean & SD] |  |  |  |
|  |  |  |  | closest SNP | most distant SNP |  |  |
|  | None | 53 |  | 0.93 ± 0.08 | 0.80 ± 0.12 |  |  |
|  | Strain only | 2 |  | 0.80 ± 0.29 | 0.84 ± 0.02 |  |  |
|  | SNP only | 54 |  | 0.87 ± 0.06 | 0.49 ± 0.26 |  |  |
|  | SNP x strain | 20 |  | 0.73 ± 0.29 | 0.46 ± 0.26 |  |  |

A further 360 of the 711 scaffolds only showed a large main effect of SNP differences, implying that they each contained at least one break in haplotype structure which differentiated the allele frequencies of its constituent SNPs, but that none of those SNPs differed significantly in frequency between the strains (Table 2A, Fig. 3C). This pattern is consistent with recombination events that predated the separation of the experimental strains and SNPs that were all unaffected by the thermal selection.

However, 83 of the 711 eligible scaffolds all yielded large main and/or interaction effects of strain, implying differentiation in at least some of their SNPs between the strains. Eight of these 83 scaffolds showed a large main effect of strain in the absence of a large SNP or interaction term (Fig. 3B). These eight are interpreted as lying within larger haplotypes containing one or more SNPs that were targeted by some form of thermal selection. The targeted SNPs (54 variants) could have been situated within the scaffold, either scored or not in our transcriptome, or beyond the scaffold boundaries and not present in our transcriptome. The remaining 75 scaffolds all involved large interaction terms, implying that some recombination had occurred in each of them in at least one, but not all strains (Fig. 3D). This strain specificity implies that the recombination occurred after the strains had been separated. The interaction also indicates that one or more SNPs in some part(s) of the scaffold also differed in frequency between strains. Most of these 75 scaffolds also involved a main effect of SNP, and two also a main effect of strain, but those main effects belie simple interpretation considering the interactions.

The 75 scaffolds with large main or interaction effects of strain included 93 of the 350 individually significant SNPs. The three scaffolds with the most significant SNPs were 252, 513 and 12047 with 11, 12, and 28 SNPs, respectively. Annotated features among the 75 scaffolds included a 9-cis-epoxycarotenoid dioxygenase (scaffold 252, with one missense and one upstream variant), a newly annotated putative E3 ubiquitin-protein ligase ORF HERC1 (scaffold 513, all 9 variants non-synonymous), the lncRNA Cproliferum_lnc.39531 (scaffold 1429, with 10 SNPs) and the NRPS_term_dom gene (scaffold 12047, with 17 downstream variants). The 93 SNPs also included all 11 of the individually significant SNPs that distinguished SS8 from SS3 and SS5 and, as noted above, all of these lay in just 5 of the 75 scaffolds.

Estimates of LD between the SNPs (mle_est) in individual scaffolds and models of their relationships to the distances between the respective SNPs could be obtained for 129 of the 711 scaffolds (Tables 2B, S5, Fig. 3). These relationships also confirmed the evidence regarding recombination that we found in the ANOVAs. Thus, there was little decline in average LD from the closest to the most distant pairs of SNPs in scaffolds not showing large main or interaction effects of SNP differences and, conversely, average LD declined markedly in scaffolds showing large main or interaction effects of SNP differences. There was no overall difference between the WT and SS strains in the extent of the LD decline among the 20 scaffolds where large interaction effects had been found (Fig. S3), although in one scaffold, 12047, the declines were more pronounced in SS3 and SS5 than SS8 and WT (Fig. 3D).

## DISCUSSION

Four years of thermal selection produced heat tolerant Symbiodiniaceae strains without a detectable contribution from new mutations in the sequenced fraction of the genome. Fewer than 1% of the alleles scored were private to a single strain, allele frequencies at 350 loci differed between strains, and the linkage patterns within scaffolds point to recombination both before and after the strains were separated. Hence, the response drew on variation that the unselected wild-type strain already carried. Benefits of standing genetic variation and recombination for rapid adaptive responses are well established in population genetic theory (27). Adaptation from standing genetic variation does not require beneficial variants to arise first, so the response rate is determined by selection alone rather than by the supply of de novo mutations followed by selection (27). Recombination acts on genetic variation to generate new combinations and to break down associations between beneficial and deleterious alleles, which has been shown experimentally to accelerate adaptation in yeast populations evolving from standing variation (28, 29). High standing genetic and clonal diversity in natural populations of diatoms, coccolithophores and dinoflagellates, suggests such responses are widespread in marine microalgae (30). Here we show that both selection and recombination among existing variants contributed to the thermal adaptation of the heat-evolved *C. proliferum* strains during experimental evolution, while de novo mutations in the sequenced scaffolds did not.

### Standing genetic variation provided the raw material for thermal adaptation

The extent of the standing variation, 15,640 SNPs in the expressed fraction alone, was surprisingly high given that the progenitor strain was established from a presumed clonal isolate picked from a single colony forming unit on solid-phase media (17). Moreover, only 6 months had elapsed between this clonal isolation and the splitting into the experimental strains (cf the four years of experimental evolution), presenting a relatively narrow window for further mutations to arise before the experimental evolution.

Three explanations could account for the higher than expected levels of polymorphism at the founding of the experimental lines. For example, the founding cell might have carried more than one copy of some or all the genome. Intraclonal genetic variation has been reported in other dinoflagellate cultures (31), and whole-genome duplication in the Symbiodiniaceae species *Durusdinium trenchii* (32), although no equivalent duplication has been reported for *C. proliferum*, which genome is treated as haploid (33). Symbiodiniaceae genomes nonetheless show high gene copy number and substantial sequence divergence between copies (14, 34). In addition, the founding cell may have been a diploid zygote formed through fusion of two separate gametes, as observed in other dinoflagellates (31, 35) and also for Symbiodiniaceae itself (36). Finally, the founding population on solid media may not have been a single cell, as a colony forming unit can arise from more than one cell. In each case the founding population would have carried polymorphism, whether spread across multiple genome copies, between two parental haplotypes, or among a small number of genetically distinct founder cells (not mutually exclusive). The fact that the SNPs found in the WT10 control strain were mostly at intermediate frequencies suggests that the number of genomes present at founding was relatively small.

Notwithstanding the small effective population size at founding, the SNPs then present occurred in a larger number of haplotype combinations. This suggests that recombination among the SNPs had occurred at some point prior to the separation of the experimental strains, and in many cases before establishment of the founding population. This could have occurred via heterothallic reproduction (i.e., fusion of gametes from two strains) or homothallic reproduction (i.e., fusion of gametes from a single clone), or both (31, 37). In *Alexandrium minutum*, planozygotes can undergo meiotic division cycles independently of resting cyst formation (38). This offers a potential mechanism for recombining existing SNPs outside the usual dormancy pathway and for how the observed genomic variation and haplotype combinations could have arisen in our strains.

### Targets of selection for enhanced thermal tolerance

A few of the 350 SNPs associated with the thermal tolerance differences between the WT and heat-evolved strains were in genes with known roles in heat-stress responses. For example, ubiquitin-related genes (18 significant SNPs) have often been linked to the processing of disordered proteins in other species exposed to heat and other stresses (39, 40). Interestingly, ubiquitin ligase activity has been flagged previously as a potential mechanism in the thermal stress response of another heat-evolved *C. proliferum* strain (SS4) (10). Other SNPs could be associated with modifications to the peroxisome architecture of the heat-evolved strains (two SNPs in peroxisome biogenesis factor 2; one SNP in peroxisomal membrane protein PMP22), which could also affect downstream peroxisome-dependent processes, such as photorespiration, beta-oxidation of fatty acids and ROS detoxification (41, 42). Additionally, the activity of 9-cis-epoxycarotenoid dioxygenase (two SNPs) has been linked to tolerance of multiple abiotic stresses in plants (43, 44), and may contribute to photoprotection and ROS scavenging in Symbiodiniaceae through changes in carotenoid metabolism (45). These SNPs might be directly involved in the enhanced thermal tolerance of the heat-evolved microalgae.

Some genes, not previously annotated or associated with thermal tolerance, contained significant SNPs. One of these (SymbC1.scaffold12047.2, 23 variants) contained the most significant SNPs. It belongs to a large nonribosomal peptide synthetase (NRPS) gene family found across Symbiodiniaceae genomes. Its precise function is not known but it shows homology to a linear gramicidin synthase subunit C with a non-ribosomal peptide synthetase terminal domain in *D. trenchii* (blastp: 75% identity, 59% coverage, e-value 0.0). The role of these candidate genes, including the NRPS gene, remains to be tested directly, for example by introducing the candidate alleles or haplotypes into a WT strain, followed by resequencing under thermal selection.

Surprisingly large proportions of both the significant and non-significant SNPs (33% and 31%, respectively) fell outside previously annotated coding regions. Some of these SNPs may lie in genes that have been missed in the reference annotation (13). Our screening of the significant SNPs previously annotated as intergenic identified 7 additional ORFs ≥ 500 nt long which BLASTp searches found matched functional genes (including an amino acid transporter on scaffold 8952), and identified 18 lncRNAs genes. Many lncRNAs have been shown to regulate gene expression in other species (46) and nearly 50,000 have been reported in various Symbiodiniaceae species, including Cladocopium, with some in the latter previously associated with gene expression during thermal stress (26). One of these (Cproliferum_lnc.39531 on scaffold 1429), in which 10 significant SNPs were found in the current study, had previously been strongly associated with the upregulation of several coding genes during heat stress (26). Thus, at least some apparently intergenic SNPs in our dataset may function in thermal stress responses and have been targets of selection in the current experiment.

All the biochemistries linked to thermal tolerance that differentiated the SS strains as a group from WT10 overlap with those identified in other experimental evolution studies of photosynthetic microorganisms. The response reported for the strains studied here occurred over a similar or smaller number of generations. The adaptation of the marine diatom *Thalassiosira pseudonana* to elevated temperature over 300 generations was associated with variants in genes related to oxidative stress responses, redox homeostasis, and transcriptional regulation (47). Similarly, the thermal adaptation of the green microalga Chlorella vulgaris after ∼100 generations involved changes in carbon metabolism, via a down-regulation of respiration compared to photosynthesis (48). In the green microalga *Picochlorum* sp. BPE23, adaptation to supra-optimal temperatures was associated with mutations in genes related to oxidative stress (e.g., a putative superoxide dismutase) with adjustments to energy metabolism and photosynthesis (49). Although the route taken by our Symbiodiniaceae strains was shaped by the standing variation available to them, and thermal tolerance may be reached by more than one genetic route, the recurrence of these broad functional categories across the four species is consistent with a common physiological basis for *in vitro* thermal adaptation in photosynthetic microalgae.

Our study has provided much less insight into the genetic basis for *in hospite* thermal tolerance. The heat-evolved SS8 strain, which conferred thermal tolerance on coral hosts (19, 23), differed from SS3 and SS5 at only 11 SNPs across five scaffolds. The specific genes implicated included a carbonic anhydrase 2, which is likely involved in carbon fixation during photosynthesis (50). While this particular gene was not differentially expressed in previous transcriptomic analyses, carbon fixation GO terms were enriched among other genes over-expressed in SS8 compared to SS3 and SS5 (19). These results are suggestive of a causal link to *in hospite* tolerance, although it is noted that our transcriptomic snapshot was taken from larvae which had not themselves been subjected to thermal stress (51). Transcriptomes of heat stressed larvae and whole genome resequencing of the strains in question would provide more insights into the genetic and epigenetic basis of enhanced *in hospite* tolerance.

### The role of recombination in enhancing thermal tolerance

Both the ANOVA and LD analyses showed patterns that are consistent with recombination within scaffolds, not only prior but also after strain separation, since commencement of experimental evolution. These events expanded the haplotype diversity available for selection to act on in each strain.

Other evidence for sexual reproduction and recombination in Symbiodiniaceae comes from genotyping of microsatellite markers in field populations (52–54) and the existence of homologues of meiosis genes in related taxa (55, 56). More direct evidence has been provided through the cytological observation of gamete conjugation, zygote formation and meiosis in approximately 1.5% of *C. latusorum* cells within *Pocillopora* corals *in situ* (36) and the observation of meiotic cells in cultures of several Symbiodiniaceae species, including the wild-type culture used in our study (57).

Some studies suggest that recombination via either homo- or heterothallism (31, 35) may be triggered by environmental stresses, such as elevated temperatures (9) and nutrient deficiencies (35, 58). Elevated expression of meiosis genes has also been recorded during dinoflagellate blooms (59), and the expression of homologues of known meiosis genes in other species has been shown to increase under elevated temperature in *C. proliferum* (9). We therefore screened the data (twenty eligible scaffolds) for evidence of reduced LD, i.e. more recombination, in the heat-evolved than WT strain. While no consistent strain differences in LD decay were found across scaffolds, the small sample size limited the power of the analyses, so the possibility that thermal stress promoted recombination in the heat-evolved strains cannot yet be discounted (see scaffold 12047 in Fig. 3D).

Given the strong LD across much of the genome, many of the significant SNPs observed could simply reflect hitchhiking (60) with closely linked genes not present in our dataset that were targeted by the selection. The LD analyses confirmed a decay of LD with physical distance between SNPs within many scaffolds, which is a well-established population genetic signature of recombination (61). Although the lack of a chromosome-level assembly prevented inter-scaffold LD analyses, the often high intra-scaffold LD suggests that some extended haplotype blocks could have exceeded the relevant scaffold boundaries. Consequently, while post-separation recombination served as a driving mechanism for generating adaptive haplotype diversity, there is still sufficient linkage to complicate the identification of individual causal mutations within these regions.

One limitation of the current study was that SNPs were called from transcriptomes and lie on scaffolds representing only 36% of the assembly length and 14% of its annotated protein-coding genes. Hence, the conclusion that new mutations contributed little applies to the sequenced fraction rather than the whole genome. A second limitation was that allele frequencies were measured in symbiont cells within coral larvae rather than in the source cultures, and each infection involved a colonization bottleneck. Random sampling at colonization would vary independently among the five or six replicate infections per strain and is accounted for in the within-strain variance, but any systematic difference among genotypes in colonization success would shift the frequencies in the same direction in every replicate. The frequencies reported here therefore may not match those in the cultures.

Notwithstanding these limitations, we concluded that the raw material for the response was available from the outset and was reorganized into new combinations during selection, so neither process had to rely on new mutations to accumulate. Our results further suggest that future experimental evolution studies aiming to develop heat-tolerant coral symbionts should start with genetically diverse populations. This could be achieved by pooling independently obtained conspecific cultures that are likely to represent distinct genotypes. Standing variation should then provide a resource which selection and recombination can draw on to increase the speed of the response.

## Supporting information

Supplementary_Material_1_SNP_table

Supplementary_Material_2_ANOVA_classification

## ACKNOWLEDGEMENTS

Funding

P.B. was supported by Macquarie University and a CSIRO Research Office Postdoctoral Fellowship. M.J.H.v.O. was supported by Australian Research Council Laureate Fellowship FL180100036.

## Author contributions

Conceptualization: P.B., J.G.O., O.R.E., M.J.H.v.O. Data analyses: P.B., J.G.O., H.L.Y.

Manuscript preparation: P.B., J.G.O., O.R.E., M.J.H.v.O., H.L.Y.

## Data and materials availability

All data needed to evaluate the conclusions in the paper are present in the paper and the Supplementary Materials. The raw sequence data are available at the NCBI Gene Expression Omnibus under accession GSE133082 and at NCBI BioProject PRJNA549921 (SRA SRP201999).

## List of Supplementary Materials

• Supplementary Methods S1 to S4

• Figs. S1 to S3

• Tables S1 to S5

• Supplementary Materials S1 and S2

