## Supplementary Methods, Figures S1-S3, and Tables S1-S5 for "Rapid thermal adaptation in coral photosymbionts draws on standing variation and recombination"

**^3^** Health & Biosecurity, Commonwealth Scientific & Industrial Research Organisation, Parkville, VIC, Australia

**^4^** Land & Water, Commonwealth Scientific & Industrial Research Organisation, Floreat, WA, Australia

**^5^** School of BioSciences, The University of Melbourne, Parkville, VIC, Australia

**^6^** Environment, Commonwealth Scientific & Industrial Research Organisation, Black Mountain, Canberra, ACT, Australia

**SUPPLEMENTARY MATERIALS**

**Supplementary Methods**

**S1. Culture origin, thermal selection and long-term maintenance**

Symbiodiniaceae cells were extracted from a colony of *Acropora tenuis* (now *A. kenti*) collected from the inshore, central Great Barrier Reef in 2010. The extracted cells were transferred into Daigo's IMK medium for marine microalgae containing antibiotics and inoculated into fresh medium monthly for five months to reduce bacterial contamination. This heterogeneous culture was subsequently plated onto IMK with antibiotics and 1% agar, and a clonal strain was established by picking cells from a single colony forming unit, which was assumed to produce a clonal culture. The resulting strain was confirmed as ITS2 type C1 (*Cladocopium proliferum*).

The clonal strain was kept at 27˚C for ~6 months, after which ten replicate subcultures of approximately the same population size (300,000 cells/mL) were subjected to a step-wise temperature increase from 27˚C to 31˚C, with each of the intervening steps at 28˚C and 30˚C lasting one month (i.e. two or three generations), and then kept at 31˚C (17).

Thermal phenotypes were characterized previously. After four years of selection (approximately 120 generations), all ten heat-evolved strains showed greater thermal resistance in culture than the wild-type control subcultures, while three conferred thermal tolerance in symbiosis with *A. kenti* larvae during a one week exposure to 31°C (19).

**S2. Sample collection and sequencing**

Transcriptome data were obtained from the ambient temperature treatment (27°C) of a previous larval infection experiment (19). Each of the coral larvae at the time of sampling harboured approximately 10,000 microalgal cells, and ten larvae for each symbiont strain were pooled into each experimental replicate. Six replicates were collected for each of SS3, SS5 and WT10, and five for SS8. The protocols for RNA extractions, library preparations and short-read 2 x 100 bp sequencing on an Illumina NovaSeq machine are described in full elsewhere (19). About 2 billion 2x100 bp paired end reads were obtained, at an average of ~87 million reads per sample (19). Only about 33% (~29 million) of the reads per sample mapped to the 1.2 Gbp reference *C. proliferum* genome but this was expected given that the sequencing included the coral host. The transcriptomes included 15,260 annotated genes with at least 20x coverage (and 22,561 genes with at least one read coverage), as compared to the 35,913 genes in the current reference genome assembly for the species (25).

**S3. SNP calling and filtering**

SNP calling was performed using the Genome Analysis Toolkit best practices workflow (GATK version 4.3.0.0, "RNAseq short variant discovery (SNPs + Indels)" at https://gatk.broadinstitute.org/hc/en-us/articles/360035531192). As recommended, RNA-seq reads were mapped with the STAR aligner (version 2.5.3a) in 2-pass mode to the *C.* *proliferum* v1.0 reference genome available at reefgenomics.org, in order to separate host from Symbiodiniaceae reads. Duplicate Symbiodiniaceae reads were then marked with Picard (version 2.6.0) and the alignments reformatted and controlled for quality with the GATK tool SplitNCigarReads. Variants were called with GATK HaplotypeCaller and calibrated with two passes of Base Quality Score Recalibration (BQSR), using variants filtered as per the criteria below from the first pass as known sites in the second pass. Following previous population genomic studies of coral holobionts (67,68), a robust filter was then applied with GATK’s VariantFiltration function to remove variants with a read depth DP score < 20, strand bias FS score > 60, variant confidence QD score < 2 or sequencing bias SOR > 3. To further minimise the influence of sequencing errors and false positives, singleton variants were also removed (--mac 2 and --min-alleles 2, using vcftools v0.1.16), as were SNPs for which sufficient reads were not available in all samples and strains (--max-missing 1.0).

**S4. Linkage disequilibrium analyses**

LDx software was used to obtain maximum likelihood estimates (mle_est) of linkage disequilibrium (LD) between pairs of SNPs in each scaffold containing at least six SNPs. Consistent with the original filters for SNP identification, the following input parameters were imposed: min read depth -l 20, max read depth -h 5000, insert size -s 200, quality score -q 20, min allele frequency -a 0.01, min intervals -I 5. Models could be obtained for 129 of the 711 scaffolds above, the major limitation being SNP numbers: models were obtained for 68% of scaffolds with ≥12 SNPs but only 16% of scaffolds with just six or seven SNPs.

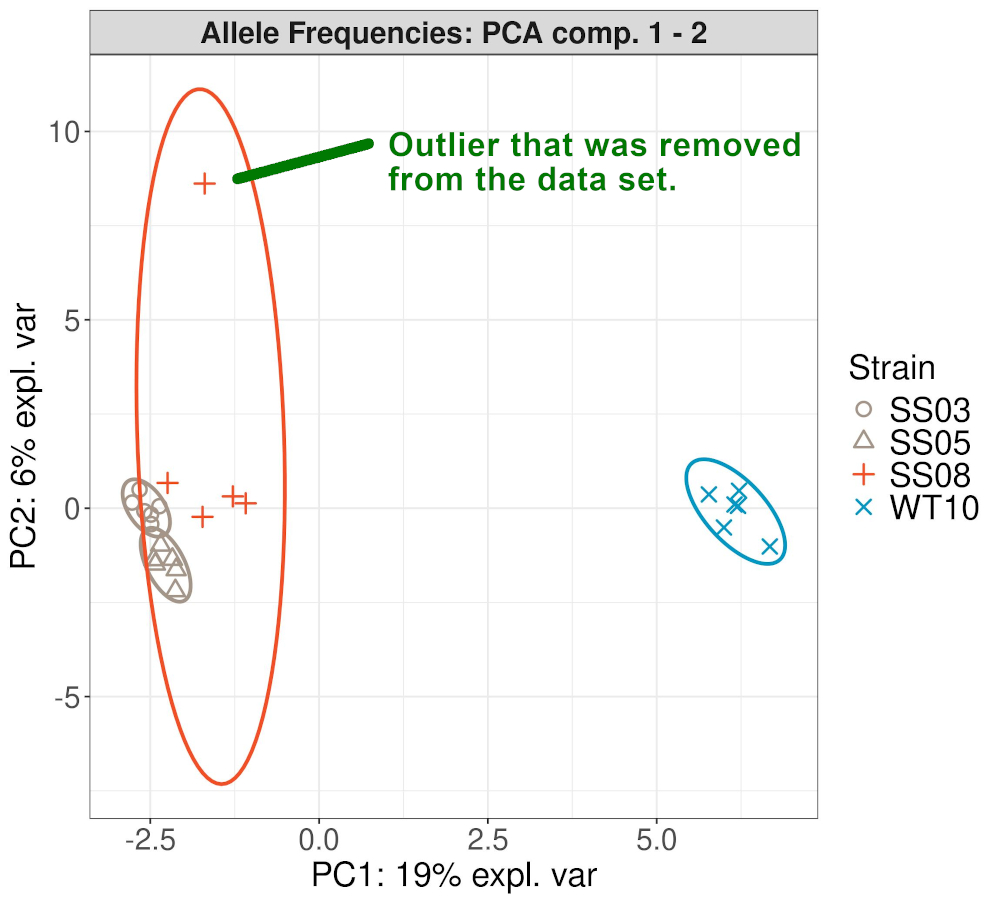

**Figure S1. Principal component analysis based on allele frequencies using microalgae SNP data of all replicates.** The plot shows that one of the SS8 replicates is an outlier and skewed data projections - hence it was removed from further downstream analyses.

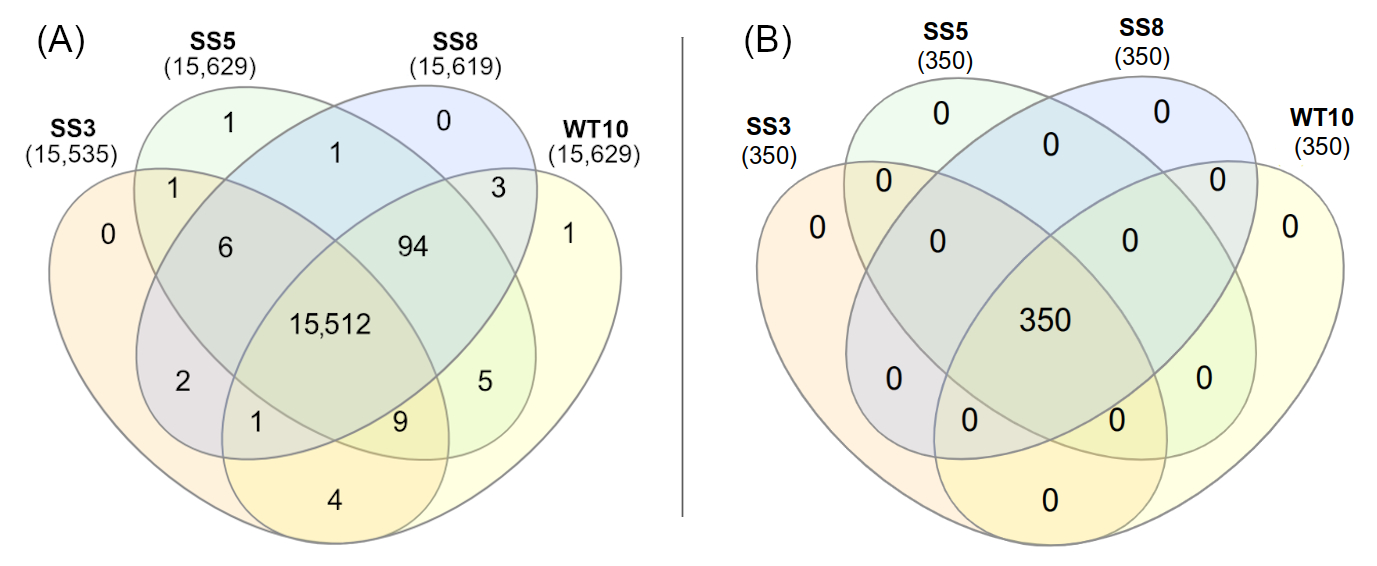

**Figure S2. Venn-diagram showing private and shared alleles (includes both fixed and polymorphic alleles).** (A) Included includes all SNPs with private reference alleles, (B) significant SNPs based on allele frequencies. Numbers in parentheses show the total SNPs for a respective strain.

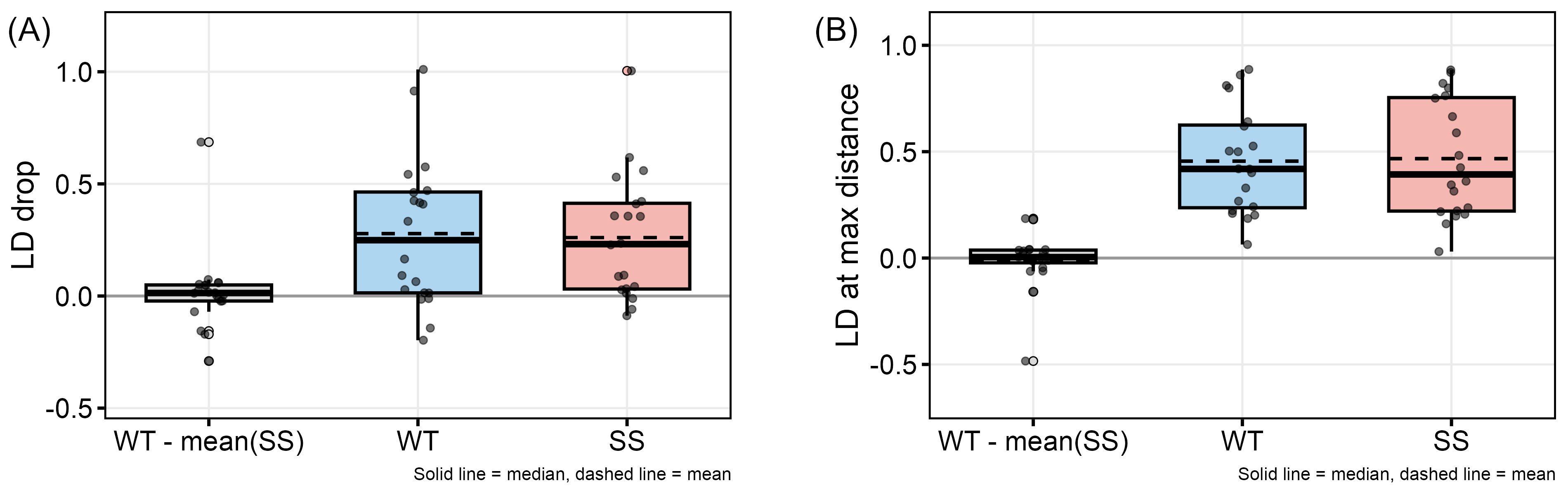

**Figure S3. Linkage disequilibrium (LD) decay comparison between wild type and heat evolved strains using scaffolds that had a significant interaction term (n = 20).** Linkage disequilibrium decay does not differ systematically between wildtype and heat-selected lineages across the interaction-significant scaffolds. Each point is one scaffold, boxes show the median (solid line), mean (dashed line), and the 25th percentile at the bottom to the 75th percentile at the top. (A) LD drop shows how much linkage disequilibrium falls off from short to long distances along a scaffold. A larger value means the linkage breaks down more, which points to more recombination. (B) LD at maximum distance shows how much linkage disequilibrium is still left at the farthest distances along a scaffold. A larger value means more linkage remains, which points to less recombination. In both panels the left box shows the difference between the respective WT and the three SS strains, and the middle and right boxes show the WT and SS values separately.

**Table S1. Overview of SNP distribution across the top scaffolds in the microalgal genome.** Shown are the top 20 scaffold with the most SNPs in descending order. The number of significant SNPs are shown, along with the scaffold length and genes on scaffold.

|  | **Scaffold ID** | SNP loci on scaffold | Significant SNPs on scaffold | Scaffold Length | Genes on Scaffold (total) |
| --- | --- | --- | --- | --- | --- |
| 1 | SymbC1.scaffold4420 | 89 | 0 | 66,563 | 4 |
| 2 | SymbC1.scaffold4221 | 72 | 0 | 69,138 | 3 |
| 3 | SymbC1.scaffold5575 | 56 | 0 | 74,883 | 5 |
| 4 | SymbC1.scaffold1202 | 44 | 0 | 146,066 | 4 |
| 5 | SymbC1.scaffold282 | 43 | 0 | 246,197 | 16 |
| 6 | SymbC1.scaffold2391 | 41 | 0 | 201,278 | 13 |
| 7 | SymbC1.scaffold13474 | 40 | 0 | 11,484 | 1 |
| 8 | SymbC1.scaffold11424 | 39 | 0 | 18,451 | 2 |
| 9 | SymbC1.scaffold2211 | 38 | 0 | 107,249 | 6 |
| 10 | SymbC1.scaffold3349 | 36 | 0 | 83,046 | 8 |
| 11 | SymbC1.scaffold16252 | 35 | 0 | 6,761 | 2 |
| 12 | SymbC1.scaffold127 | 34 | 0 | 308,225 | 15 |
| 13 | SymbC1.scaffold5749 | 34 | 0 | 51,883 | 0 |
| 14 | SymbC1.scaffold13933 | 34 | 0 | 10,589 | 1 |
| 15 | SymbC1.scaffold451 | 34 | 0 | 212,229 | 10 |
| 16 | SymbC1.scaffold2508 | 34 | 0 | 100,434 | 5 |
| 17 | SymbC1.scaffold2502 | 33 | 0 | 108,143 | 6 |
| 18 | SymbC1.scaffold1235 | 33 | 0 | 152,160 | 9 |
| 19 | SymbC1.scaffold16794 | 32 | 0 | 6,265 | 0 |
| 20 | SymbC1.scaffold14903 | 31 | 1 | 8,750 | 0 |

**Table S2. Summary of location and genes of 350 significant SNPs.** The scaffolds are sorted in descending order with respect to the number of significant SNPs.

| **Scaffold ID** | **SNPs on scaffold** | **Significant SNPs on scaffold** | **Scaffold length** | **Genes on scaffold (total)** |
| --- | --- | --- | --- | --- |
| SymbC1.scaffold12047 | 28 | 23 | 16,330 | 1 |
| SymbC1.scaffold1429 | 20 | 10 | 134,713 | 1 |
| SymbC1.scaffold513 | 12 | 9 | 202,056 | 6 |
| SymbC1.scaffold1521 | 10 | 9 | 233,770 | 17 |
| SymbC1.scaffold252 | 11 | 8 | 253,846 | 21 |
| SymbC1.scaffold2942 | 8 | 8 | 90,396 | 6 |
| SymbC1.scaffold5652 | 8 | 8 | 52,707 | 4 |
| SymbC1.scaffold3511 | 7 | 7 | 79,837 | 0 |
| SymbC1.scaffold4378 | 7 | 7 | 67,137 | 4 |
| SymbC1.scaffold3949 | 15 | 7 | 72,764 | 4 |
| SymbC1.scaffold15669 | 7 | 7 | 18,047 | 2 |
| SymbC1.scaffold5104 | 6 | 6 | 105,438 | 1 |
| SymbC1.scaffold2004 | 6 | 6 | 218,015 | 10 |
| SymbC1.scaffold14983 | 6 | 6 | 8,632 | 1 |
| SymbC1.scaffold4829 | 7 | 6 | 61,765 | 2 |
| SymbC1.scaffold4052 | 5 | 5 | 85,282 | 1 |
| SymbC1.scaffold9398 | 5 | 5 | 26,510 | 3 |
| SymbC1.scaffold12934 | 5 | 5 | 15,183 | 2 |
| SymbC1.scaffold936 | 4 | 4 | 163,011 | 6 |
| SymbC1.scaffold1415 | 4 | 4 | 134,870 | 11 |
| SymbC1.scaffold235 | 10 | 4 | 258,262 | 15 |
| SymbC1.scaffold3092 | 4 | 4 | 87,357 | 5 |
| SymbC1.scaffold8674 | 4 | 4 | 30,413 | 1 |
| SymbC1.scaffold3829 | 31 | 4 | 74,439 | 2 |
| SymbC1.scaffold7127 | 4 | 4 | 40,657 | 1 |
| SymbC1.scaffold8952 | 26 | 4 | 52,236 | 2 |
| SymbC1.scaffold1548 | 4 | 4 | 128,746 | 5 |
| SymbC1.scaffold2220 | 3 | 3 | 106,778 | 1 |
| SymbC1.scaffold5135 | 3 | 3 | 103,774 | 0 |
| SymbC1.scaffold1717 | 3 | 3 | 121,835 | 4 |
| SymbC1.scaffold1154 | 22 | 3 | 221,547 | 5 |
| SymbC1.scaffold580 | 3 | 3 | 194,049 | 7 |
| SymbC1.scaffold4974 | 6 | 3 | 60,039 | 5 |
| SymbC1.scaffold75 | 3 | 3 | 363,556 | 10 |
| SymbC1.scaffold7281 | 4 | 3 | 77,729 | 3 |
| SymbC1.scaffold304 | 4 | 3 | 238,993 | 9 |
| SymbC1.scaffold5401 | 5 | 3 | 55,333 | 3 |
| SymbC1.scaffold2331 | 3 | 3 | 104,324 | 4 |
| SymbC1.scaffold2087 | 2 | 2 | 110,959 | 5 |
| SymbC1.scaffold10530 | 2 | 2 | 35,205 | 0 |
| SymbC1.scaffold5242 | 2 | 2 | 57,005 | 2 |
| SymbC1.scaffold4589 | 2 | 2 | 116,230 | 2 |
| SymbC1.scaffold2613 | 22 | 2 | 198,993 | 8 |
| SymbC1.scaffold3685 | 2 | 2 | 78,998 | 4 |
| SymbC1.scaffold1720 | 2 | 2 | 125,808 | 2 |
| SymbC1.scaffold2074 | 8 | 2 | 118,836 | 3 |
| SymbC1.scaffold5600 | 2 | 2 | 53,301 | 4 |
| SymbC1.scaffold1161 | 6 | 2 | 148,538 | 7 |
| SymbC1.scaffold1282 | 19 | 2 | 142,097 | 0 |
| SymbC1.scaffold1044 | 2 | 2 | 156,110 | 6 |
| SymbC1.scaffold4879 | 2 | 2 | 61,040 | 3 |
| SymbC1.scaffold518 | 3 | 2 | 298,453 | 8 |
| SymbC1.scaffold4064 | 2 | 2 | 71,493 | 0 |
| SymbC1.scaffold5416 | 3 | 2 | 55,149 | 3 |
| SymbC1.scaffold5049 | 2 | 2 | 59,338 | 7 |
| SymbC1.scaffold1117 | 3 | 2 | 154,412 | 1 |
| SymbC1.scaffold196 | 4 | 2 | 272,030 | 6 |
| SymbC1.scaffold9812 | 2 | 2 | 28,257 | 1 |
| SymbC1.scaffold279 | 3 | 2 | 245,894 | 7 |
| SymbC1.scaffold1937 | 11 | 2 | 115,043 | 3 |
| SymbC1.scaffold3111 | 5 | 1 | 87,159 | 5 |
| SymbC1.scaffold1827 | 1 | 1 | 118,359 | 3 |
| SymbC1.scaffold324 | 4 | 1 | 379,327 | 7 |
| SymbC1.scaffold2928 | 1 | 1 | 91,131 | 3 |
| SymbC1.scaffold4411 | 1 | 1 | 67,055 | 2 |
| SymbC1.scaffold5935 | 1 | 1 | 50,027 | 5 |
| SymbC1.scaffold26245 | 1 | 1 | 1,909 | 1 |
| SymbC1.scaffold13086 | 8 | 1 | 12,507 | 0 |
| SymbC1.scaffold1842 | 1 | 1 | 118,211 | 8 |
| SymbC1.scaffold1708 | 1 | 1 | 122,785 | 4 |
| SymbC1.scaffold3500 | 3 | 1 | 119,578 | 7 |
| SymbC1.scaffold4106 | 1 | 1 | 70,627 | 3 |
| SymbC1.scaffold3612 | 1 | 1 | 78,539 | 7 |
| SymbC1.scaffold1314 | 2 | 1 | 140,127 | 0 |
| SymbC1.scaffold1267 | 1 | 1 | 142,250 | 6 |
| SymbC1.scaffold1500 | 1 | 1 | 162,292 | 2 |
| SymbC1.scaffold5744 | 1 | 1 | 52,011 | 1 |
| SymbC1.scaffold167 | 7 | 1 | 451,144 | 10 |
| SymbC1.scaffold13187 | 1 | 1 | 12,156 | 2 |
| SymbC1.scaffold5414 | 1 | 1 | 55,159 | 2 |
| SymbC1.scaffold473 | 1 | 1 | 208,549 | 5 |
| SymbC1.scaffold13211 | 1 | 1 | 12,192 | 0 |
| SymbC1.scaffold2987 | 1 | 1 | 89,862 | 3 |
| SymbC1.scaffold17803 | 1 | 1 | 4,974 | 1 |
| SymbC1.scaffold3220 | 1 | 1 | 84,951 | 2 |
| SymbC1.scaffold2962 | 1 | 1 | 90,278 | 6 |
| SymbC1.scaffold4657 | 1 | 1 | 63,705 | 2 |
| SymbC1.scaffold6597 | 1 | 1 | 44,520 | 0 |
| SymbC1.scaffold14903 | 31 | 1 | 8,750 | 0 |
| SymbC1.scaffold3812 | 1 | 1 | 74,950 | 8 |
| SymbC1.scaffold2299 | 1 | 1 | 104,939 | 4 |
| SymbC1.scaffold25965 | 1 | 1 | 1,983 | 0 |
| SymbC1.scaffold198 | 3 | 1 | 270,853 | 10 |
| SymbC1.scaffold5888 | 1 | 1 | 50,409 | 5 |
| SymbC1.scaffold7250 | 1 | 1 | 39,702 | 1 |
| SymbC1.scaffold82 | 1 | 1 | 357,611 | 4 |
| SymbC1.scaffold927 | 1 | 1 | 163,458 | 5 |
| SymbC1.scaffold860 | 12 | 1 | 168,629 | 9 |
| SymbC1.scaffold35833 | 2 | 1 | 1,196 | 0 |
| SymbC1.scaffold2404 | 1 | 1 | 111,964 | 5 |
| SymbC1.scaffold655 | 1 | 1 | 186,312 | 1 |
| SymbC1.scaffold8147 | 3 | 1 | 33,866 | 1 |
| SymbC1.scaffold1816 | 1 | 1 | 126,013 | 6 |
| SymbC1.scaffold2401 | 3 | 1 | 102,948 | 4 |
| SymbC1.scaffold6543 | 1 | 1 | 51,681 | 1 |
| SymbC1.scaffold11402 | 1 | 1 | 35,065 | 1 |
| SymbC1.scaffold9385 | 1 | 1 | 26,674 | 3 |
| SymbC1.scaffold4266 | 1 | 1 | 68,647 | 0 |
| SymbC1.scaffold6722 | 15 | 1 | 92,680 | 3 |
| SymbC1.scaffold1921 | 1 | 1 | 115,427 | 7 |
| SymbC1.scaffold13025 | 1 | 1 | 12,700 | 0 |
| SymbC1.scaffold2858 | 2 | 1 | 92,725 | 7 |
| SymbC1.scaffold7985 | 1 | 1 | 34,950 | 0 |
| SymbC1.scaffold3664 | 1 | 1 | 77,346 | 7 |
| SymbC1.scaffold2710 | 1 | 1 | 95,803 | 4 |
| SymbC1.scaffold15906 | 1 | 1 | 7,260 | 1 |
| SymbC1.scaffold6697 | 2 | 1 | 43,619 | 2 |
| SymbC1.scaffold10186 | 8 | 1 | 22,991 | 2 |
| SymbC1.scaffold581 | 1 | 1 | 193,622 | 7 |
| SymbC1.scaffold2438 | 1 | 1 | 102,263 | 3 |
| SymbC1.scaffold2445 | 1 | 1 | 102,383 | 2 |
| SymbC1.scaffold965 | 1 | 1 | 161,537 | 10 |
| SymbC1.scaffold700 | 5 | 1 | 181,746 | 10 |
| SymbC1.scaffold1181 | 1 | 1 | 147,007 | 4 |
| SymbC1.scaffold27164 | 1 | 1 | 1,794 | 0 |
| SymbC1.scaffold12498 | 1 | 1 | 14,205 | 0 |
| SymbC1.scaffold3873 | 3 | 1 | 74,019 | 3 |
| SymbC1.scaffold4486 | 1 | 1 | 65,991 | 2 |
| SymbC1.scaffold4514 | 1 | 1 | 65,386 | 3 |
| SymbC1.scaffold4906 | 1 | 1 | 60,655 | 1 |
| SymbC1.scaffold1035 | 2 | 1 | 164,043 | 1 |
| SymbC1.scaffold9698 | 1 | 1 | 25,182 | 1 |
| SymbC1.scaffold761 | 1 | 1 | 303,678 | 6 |
| SymbC1.scaffold7607 | 7 | 1 | 37,144 | 5 |
| SymbC1.scaffold3139 | 1 | 1 | 86,956 | 2 |
| SymbC1.scaffold14538 | 4 | 1 | 10,359 | 0 |
| SymbC1.scaffold1426 | 13 | 1 | 142,247 | 7 |
| SymbC1.scaffold5169 | 1 | 1 | 57,560 | 2 |
| SymbC1.scaffold6443 | 1 | 1 | 45,773 | 0 |
| SymbC1.scaffold1204 | 2 | 1 | 165,850 | 4 |
| SymbC1.scaffold8467 | 1 | 1 | 31,784 | 3 |
| SymbC1.scaffold7176 | 8 | 1 | 45,723 | 4 |
| SymbC1.scaffold6694 | 2 | 1 | 50,729 | 2 |
| SymbC1.scaffold3649 | 1 | 1 | 77,554 | 2 |
| SymbC1.scaffold792 | 1 | 1 | 174,056 | 3 |
| SymbC1.scaffold919 | 1 | 1 | 164,124 | 3 |
| SymbC1.scaffold137 | 4 | 1 | 300,232 | 2 |
| SymbC1.scaffold1514 | 1 | 1 | 130,156 | 4 |
| SymbC1.scaffold1914 | 2 | 1 | 116,395 | 6 |
| SymbC1.scaffold210 | 28 | 1 | 267,022 | 13 |
| SymbC1.scaffold2353 | 4 | 1 | 103,797 | 3 |
| SymbC1.scaffold1882 | 1 | 1 | 116,908 | 2 |
| SymbC1.scaffold10275 | 1 | 1 | 22,414 | 1 |
| SymbC1.scaffold6798 | 1 | 1 | 42,941 | 1 |
| SymbC1.scaffold9048 | 1 | 1 | 28,505 | 2 |
| **TOTAL: 155 scaffolds** | **689 SNPs** | **350 sign. SNPs** | **16,457,840 bp** | **589 genes** |

**Table S3** $\chi^{2}$**tests** comparing the allele frequency distributions of the 350 significant SNPs among the respective strains.

| **comparison** | **X_squared** | **df** | **p_value** | **p_adjusted** | **significant** |
| --- | --- | --- | --- | --- | --- |
| WT vs SS3 | 434.021 | 19 | 3.58E-80 | 1.79E-79 | *** |
| WT vs SS5 | 414.469 | 19 | 4.27E-76 | 2.14E-75 | *** |
| WT vs SS8 | 447.162 | 19 | 6.46E-83 | 3.23E-82 | *** |
| SS3 vs SS5 | NaN | 19 | NaN | NaN | Not sign. |
| SS3 vs SS8 | 221.578 | 19 | 1.66E-36 | 8.32E-36 | *** |
| SS5 vs SS8 | 224.507 | 19 | 4.29E-37 | 2.15E-36 | *** |

**Table S4. Summary of selected functional annotations that impacted by the 350 significant SNPs.** Top of the table is showing the comparison WT vs SS, while the comparison SS8 vs SS3+SS5 is shown on the bottom.

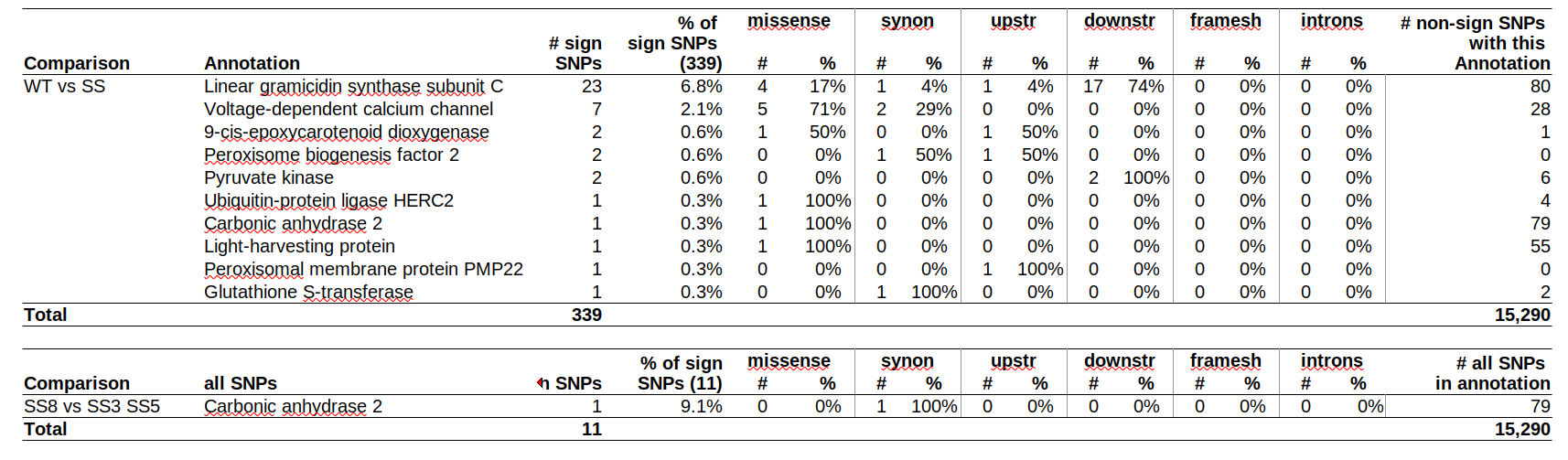

**Table S5. Summary of scaffold with linkage disequilibrium data.** LD on scaffolds was categorized in category 1) declining LD, 2) LD not changing, 3) LD showing differences between strains.

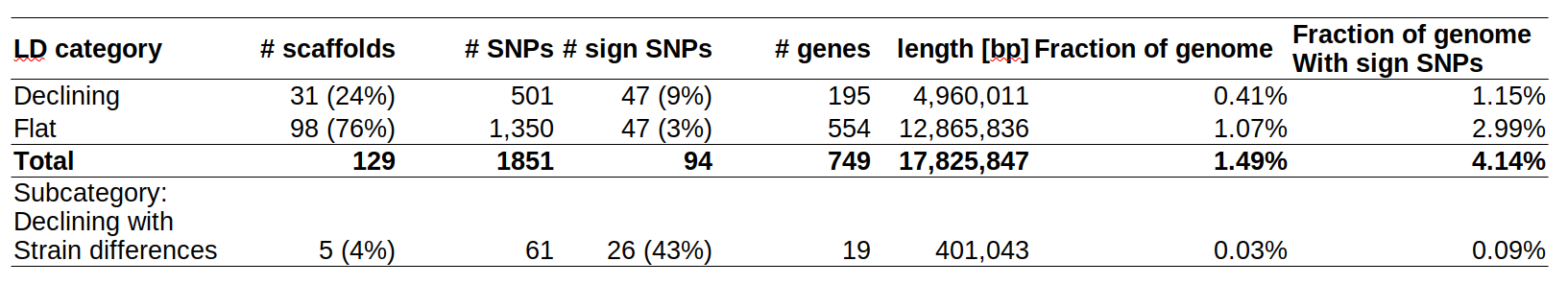
